# Exploring GPCR-mediated optogenetic modulation of seizure network in a pig model of Temporal Lobe Epilepsy

**DOI:** 10.1101/2025.01.23.634545

**Authors:** Su-youne Chang, Karni Lev Bar-Or, Fillip Mivalt, Daniela Maltais, Yoav Kfir, Jiwon Kim, Inyong Kim, Matan Hershko, Jenn Daily, Evyatar Swissa, Ofir Levi, Sasha Devore, Ofer Yizhar, Yotam Eldar, Gregory A. Worrell

## Abstract

**Rationale:** Optogenetics offers unmatched cellular specificity and control over cellular activity. Various opsins have been tested in animal models of epilepsy, each contributing to our understanding of seizure circuit dynamics. However, inhibitory optogenetic tools based on microbial rhodopsins have low light sensitivity and, thus, are less suitable for applications involving larger brains. We evaluated eOPN3, a red-shifted, highly sensitive inhibitory G-protein coupled receptor opsin in a porcine seizure model using integrated electro-optical sensing and modulation. The results demonstrated the feasibility of eOPN3 circuit modulation in a large animal epilepsy model.

**Methods:** MRI-guided stereotactic surgery was used to deliver 20–60 µL of AAV-eOPN3 (AAV5-/AAV9-CaMKII-eOPN3-mScarlet) into the hippocampus (HPC) of three Göttingen minipigs. Each hemisphere received either an active or a control viral vector (AAV5/9-CaMKII-mScarlet) with gadolinium to visualize the injection sites and diffusion volume via post-operative MRI. Two to three months post-injection, bilateral deep brain stimulation electrodes integrated with optic fibers were stereotactically implanted into the anterior nucleus of the thalamus (ANT) and HPC to assess: 1) opsin expression using fiber photometry, 2) optogenetic modulation of stimulation evoked response potentials (SERPs), 3) induction and propagation of seizure-like activity via intrahippocampal kainic acid (KA) injection, and 4) optogenetic modulation of KA-induced seizure activity. After the electrophysiology recording, brains were harvested for histological analysis to evaluate injection target precision, eOPN3 expression, and estimate eOPN3-modulated volume.

**Results:** eOPN3 expression was confirmed during surgery via fiber photometry. ANT electrical stimulation elicited robust SERPs in the HPCs, which were attenuated by HPC light illumination. HPC stimulation similarly induced SERPs in the ipsilateral ANT and the contralateral HPC. The HPC stimulation-induced SERPs were significantly reduced by illuminating the site of the recording areas, the ipsilateral ANT and the contralateral HPC, demonstrating the optogenetic inhibition of the synaptic release from the HPC. KA injection into the HPC induced 20-30 Hz seizure-like activity. The ANT and HPC light illumination suppressed the localized KA-induced seizure activity in the early stage. However, after the generalization of KA-induced seizures, the ANT-HPC illumination lost efficacy for the control of seizures. Histological analysis confirmed eOPN3 expression in the HPC, ANT and other Papez circuit nodes.

**Conclusion:** Our pilot study highlights that eOPN3-mediated inhibition alters SERP and the latency and spread of KA-induced seizure-like activity. We developed a platform incorporating pre- and postoperative MRI for precise viral vector delivery, real-time fiber photometry for quantifying opsin expression, and integrated electro-optical sensing and stimulation to assess optogenetic efficacy in a large animal model. The large animal model provides a solid foundation for future translational research to develop electro-optical devices and cellular therapies for human epilepsy.

## Introduction

Epilepsy, a chronic neurological disorder, is characterized by recurrent seizures and comorbidities associated with neural network dysfunction arising from an imbalance between excitatory and inhibitory neuronal activity^1-4^. Among its various forms, mesial temporal lobe epilepsy (mTLE) poses significant challenges due to its resistance to pharmacological treatments ^2,5,6,7,8^. Targeted circuit-based therapies have thus emerged as a promising avenue to restore neural homeostasis, alleviate seizures, and improve cognitive and psychiatric comorbidities^9-12^. However, currently available neuromodulation therapies, including deep brain stimulation (DBS), responsive neurostimulation (RNS), and vagus nerve stimulation (VNS), lack cell-specificity and utilize nonspecific modulation of various neuron types.

Optogenetics employs light to control genetically defined populations of neurons and has transformed neuroscience by enabling precise spatiotemporal manipulation of cell-type specific neural activity^13-15^. Current investigational therapies involve using optogenetic tools to restore vision in patients with retinal degeneration by targeting surviving retinal cells with light-sensitive proteins^16^. Early clinical trials have demonstrated the feasibility and safety of these approaches, offering proof-of-concept for optogenetic applications in human therapeutic contexts^16,17^. These advancements underscore the potential for expanding optogenetic techniques beyond retinal disorders to other neurological conditions, including epilepsy.

In epilepsy, optogenetic interventions offer the potential to restore the excitation-inhibition balance by selectively activating inhibitory interneurons or suppressing hyperactive excitatory neurons. These theoretical optogenetic treatment strategies have been proven successful in rodent models. Among others, Paz et. al.^18^ showed that stimulating halorhodopsin-expressing thalamocortical neurons with yellow light instantaneously aborts seizures in a rat model of photothrombotic cortical stroke; and depolarizing thalamocortical neurons expressing stabilized step function opsins stops seizures in rat and mouse models of generalized epilepsy^19^. Krook-Magnuson et al. used a closed-loop paradigm to show that either inhibiting halorhodopsin-expressing hippocampal excitatory neurons or activating channel rhodopsin-2 expressing GABAergic neurons stops seizures in a pilocarpine-induced mouse model of temporal lobe epilepsy^20^. These pioneering rodent studies revealed the promise of optogenetic strategies to treat seizures. However, translating these findings to human applications remains a considerable challenge, limited by the constraints of current experimental models, implantable device technology, opsin’s light sensitivity required for activation, illumination technology, long-term biocompatibility, and safety.

To address these challenges, organotypic hippocampal slice cultures derived from human tissue provide a versatile experimental platform to test and refine optogenetic approaches, enabling the detailed study of human-specific neuronal responses to viral delivery and network dynamics under controlled conditions^21-23^. Recent research has demonstrated that optogenetic manipulation of excitability in human hippocampal slice cultures can effectively modulate network activity and provides some initial evidence for the feasibility of human applications^24^. Despite these contributions, such models face inherent limitations, including short-term viability, limited circuit complexity, and discrepancies between in vitro and in vivo conditions. Lastly, rodent and human tissue culture models do not address the real-world challenges of developing an implantable electro-optical system for detecting seizures and controlling epileptogenic circuits in human epilepsy.

To begin to address these challenges, large animal models such as pigs, can provide a robust platform to develop the opsins, devices, and technology designed for human application. Pigs possess a gyrencephalic brain with anatomical and physiological features closely resembling those of humans, making them highly suitable for translational neuroscience research^25-28^. Their larger brain size supports surgical interventions and advanced imaging, enabling comprehensive evaluations of therapeutic strategies commonly used in human epilepsy. In epilepsy research, pig models can uniquely facilitate the long-term study of optogenetic interventions on complex neural circuits, addressing safety and efficacy concerns of opsins and devices designed for human application.

In this study, we develop a large animal platform to evaluate optogenetic interventions for mTLE. Using pig models, we sought to overcome the limitations of in vitro, ex vivo, and small animal studies. Our approach focused on inhibiting glutamatergic hippocampal neurons by expressing eOPN3, an inhibitory, bistable GPCR opsin, predominantly in these cells. eOPN3 suppresses synaptic transmission via the G_i/o_ signaling pathway (a natural, endogenous inhibitory mechanism) and features a broad action spectrum extending into the red-light range (∼630 nm) with low-intensity illumination requirements^29^. These properties make eOPN3 a promising tool for translational applications. By employing this optogenetic approach in a large animal model, we aim to advance the potential of therapies for drug-resistant epilepsy and contribute to developing more effective treatment strategies.

### Materials and Methods Subjects and Animal Care

All study procedures adhered to the National Institutes of Health Guidelines for Animal Research (Guide for the Care and Use of Laboratory Animals) and were approved by the Mayo Clinic Institutional Animal Care and Use Committee. Three healthy female Gottingen minipigs (n = 3), weighing 25 ± 5 kg at the time of surgery, were used for the study. Post-surgery, the animals were housed individually under controlled conditions (45% humidity, 21°C temperature) with ad libitum access to water and once-daily feeding. Upon reaching a weight of >40 kg, the diet was modified to include predominantly oats to manage weight gain until euthanasia. AAVs encoding eOPN3 or mScarlet were produced by and are available from the Viral Vector Core at the University of Zurich (https://vvf.ethz.ch/).

### MRI-Guided AAV-opsin Injection Surgery

Subjects were pre-medicated with Telazol (5 mg/kg, intramuscular) and Xylazine (2 mg/kg, intramuscular), followed by endotracheal intubation and intravenous catheterization. Anesthesia was maintained with 1.5–3% isoflurane, and vital signs, including heart rate (∼120 bpm), body temperature (36–37°C), and oxygen saturation, were continuously monitored. Respiratory rates were stabilized at 12 breaths per minute.

A Leksell stereotactic system (Elekta, Stockholm, Sweden) adapted for swine was employed alongside a 3.0-T MR scanner (Signa HDx, General Electric) equipped with a custom 4-channel phased-array RF coil. Brainlab software was used for stereotactic targeting based on MR images and a pig brain atlas. Injection sites in the anterior nucleus of the thalamus (ANT) and hippocampus (HPC) were determined.

After aseptic preparation, burr holes (∼1–2 mm) were drilled above the target areas. A Hamilton syringe attached to a micropump delivered viral suspensions (1.0–2.0 μl/min) containing gadobutrol (Gd, 0.5% v/v) to confirm injection accuracy and diffusion. For eOPN3 expression, ssAAV-5/2-mCaMKIIα(short)-eOPN3_mScarlet-WPRE-synp(A) and ssAAV-9/2-mCaMKIIα(short)-eOPN3_mScarlet-WPRE-synp(A) were used, and the viral stock solutions were diluted to 2 × 10^12^ vg/ml for the injection. The injection site, volume, and subject information are shown in Table 1.

**Table 1.** Subject and experimental information. Three pigs were used in the study. The viral vector serotype, injection sites, volume, opsin type, and tested experiments are shown (R; right hemisphere, L; left hemisphere, ANT; anterior nucleus of the thalamus, HPC; hippocampus, LFP; local field potential, SERP; stimulation-evoked response potential).

|  | Pig 1 |  |  | Pig 2 |  |  | Pig 3 |  |
| --- | --- | --- | --- | --- | --- | --- | --- | --- |
| AAV Serotype | AAV5/2 |  |  | AAV9/2 |  |  | AAV9/2 |  |
| Injection site | R ANT | L HPC | R HPC | R ANT | L HPC | R HPC | L HPC | R HPC |
| Injection volume | 20 μl | 20 μl X 3 | 20 μl X 3 | 20 μl | 20 μl X 3 | 20 μl X 3 | 20 μl | 20 μl X 3 |
| Expressed Opsin | Control | eOPN3 | Control | Control | eOPN3 | Control | eOPN3 | eOPN3 |
| Analysis | LFP, SERP, and Histology |  |  | LFP, SERP, and Histology |  |  | Histology |  |

A post-operative MRI was performed to confirm injection accuracy. Animals were recovered and maintained for up to three months before subsequent procedures. Sutures were removed 10– 12 days after surgery.

### Imaging for Surgery

MRI-guided implantation was performed using the same 3.0-T MR scanner. Magnetization-Prepared Rapid Gradient Echo (MPRAGE) and Fast Gray Matter Acquisition T1 Inversion Recovery (FGATIR) sequences provided high gray-white matter contrast. Image data were integrated into a surgical navigation system (Brainlab) for precise targeting. Stereotactic coordinates were determined for DBS electrodes and optic fibers targeting the bilateral ANT and HPC.

### Optogenetic stimulation and fiber-photometry

For light delivery and fiber-photometry measurements, we used custom-made polymer fibers (1000uM, na=0.63, Prizmatix, Israel), that were sanded to generate a uniform illumination profile around the last 10 mm of fiber while the remaining illumination power from the tip of the fiber is <15% (Supplementary Figure 1B). For green and red light delivery, we used high-power Dual-Optogenetics-LED (Prizmatix, Israel) with green (530-600nm, max. power 500mW, 15mW/mm^2^ on average through the fiber) and red (610-630nm, max. power 240mW, 7mW/mm^2^ on average through the fiber). To perform fiber-photometry measurements, we used the ilFMC-G3 Mini Cube fiber photometry system (Doric Lenses, Canada) to validate mScarlet expression and optimize the fiber and electrode positioning for electrophysiological recordings.

### Intra-Operative In Vivo Electrophysiology to Validate Opsin Function

Pre-operative imaging and stereotactic navigation mirrored the AAV injection procedure. Local field potentials (LFPs) and stimulation-evoked response potentials (SERPs) were recorded through quadripolar DBS electrodes (Model 3389, Medtronic) using the Navia Workstation (Cadence Neuroscience Inc., Seattle, WA) under ketamine/xylazine anesthesia. Signals were sampled at 8 kHz and bandpass filtered between 0.01 and 1000 Hz. Continuous recordings were obtained from DBS electrodes bilaterally implanted in the ANT and HPC.

Each electrode was integrated with a single optic fiber (Supplementary Figure 1B). Opsin functionality was assessed by delivering green light (peak activity 561-nm) during single-pulse stimulation (6 mA, 250 μs, 0.1 Hz). Kainic acid (KA; 1 μg/μl) was infused (0.2–1.0 μl/min) into the HPC to induce seizures, with optogenetic intervention (described in Optogenetic stimulation and fiber-photometry) applied during observed seizure activity. After the experiment, animals were euthanized, and brains were collected for analysis.

### Analysis of Electrophysiological Data

SERPs were recorded under baseline, illumination, and recovery conditions. Responses were quantified by peak amplitude and normalized to baseline. Spectrograms of KA-induced activity were processed with a 2D Gaussian filter for visualization.

### Quantification of Transfection Efficiency

Brains were fixed in 4% paraformaldehyde, cryoprotected, and sectioned into 60-μm slices. To evaluate the expression of eOPN3 in the HPC, we performed epi-fluorescent imaging on HPC slices using a Keyence microscope. The HPC region was first identified using low magnification (2x) brightfield imaging, which provided an overview of the anatomical structure and facilitated region localization (Figure 5A). The microscope was then configured for mScarlet visualization (∼561 nm excitation, ∼586 nm emission), enabling the detection of eOPN3 expression across a wide field of view (Figure 5B). Higher magnification imaging (20x) was subsequently performed to confirm eOPN3 expression at the cellular level (Figure 5C). Fluorescent mScarlet-expressing cells were imaged and quantified using ImageJ. Transfection efficiency was calculated as the percentage of mScarlet-positive areas relative to the total HPC and pyramidal cell layers.

### Statistical Analysis

Data analysis methods were described in each result section with significance set at p < 0.05. Results are presented as mean ± SEM.

## Results

### Experimental Workflow of Optogenetic Evaluation in Pigs

The experimental design involved a two-stage surgical protocol (Figure 1): (1) a chronic procedure for viral vector delivery and (2) an acute, non-survival procedure to characterize electro-optogenetic neuromodulation within the Papez circuit.

**Figure 1.**
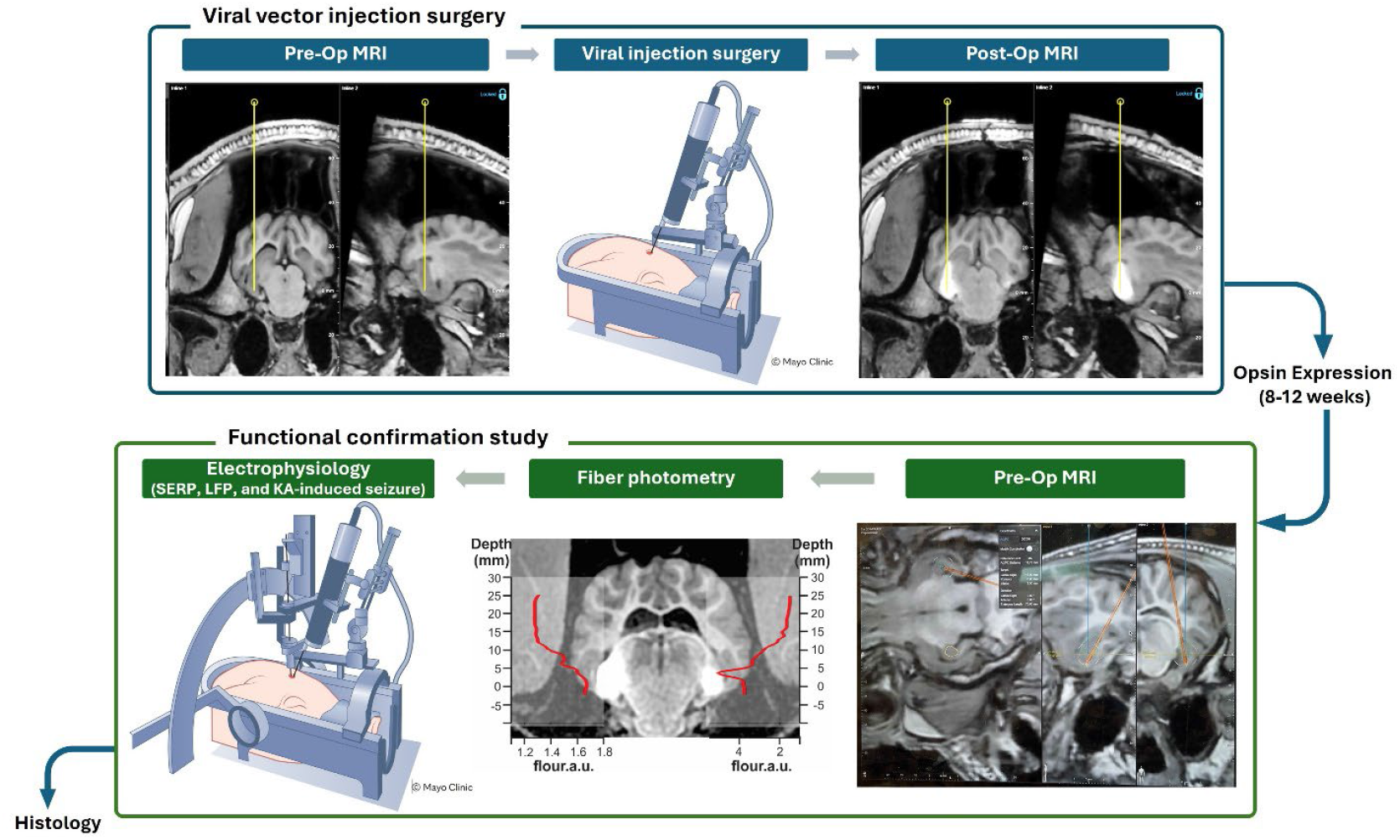
Experimental Workflow for Electro-Optogenetic Modulation in the Papez Circuit of an Acute Porcine TLE Model. The workflow involved two surgeries: (1) viral vector injection into the HPC guided by MRI and (2) an acute, non-survival procedure to evaluate electro-optogenetic neuromodulation. During the injection surgery, the AAV suspension was mixed with Gd for MRI visualization, and its target and diffusion were confirmed post-operatively. During the evaluation of acute surgery, fiber photometry confirmed eOPN3 expression, and DBS electrodes combined with optical fibers were bilaterally implanted. Neuromodulatory effects were assessed through electrical and optical stimulation with simultaneous LFP recording. Brains were collected post-experiment for histological analysis.

Accurate targeting is essential for successful neuromodulation. To facilitate this, a pre-operative MRI was conducted on the day of the surgery, utilizing MPRAGE and FGATIR sequences. These imaging modalities are routinely employed for electrode placement planning in human DBS surgeries due to their superior gray-white matter contrast^30,31^ (Figure 2A).

**Figure 2.**
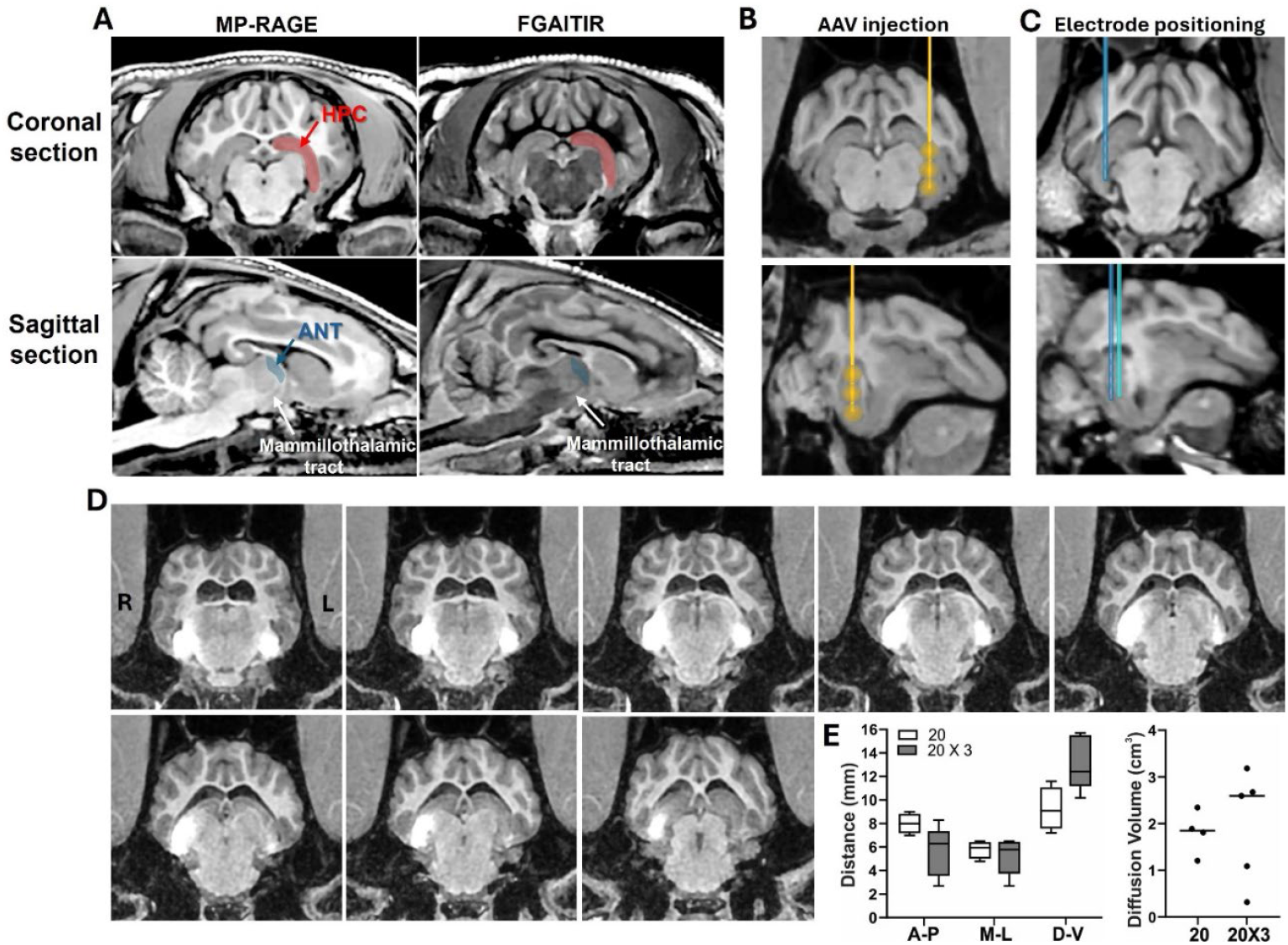
MRI-Guided Targeting for AAV Injection, Electrode Implantation, and Evaluation of Injection Success and Diffusion Volume. (A) MP-RAGE and FGAITIR images were used to identify the HPC and ANT. The mammillothalamic tract served as a landmark for delineating the ANT border. Anatomical target localization of the HPC for (B) AAV injection and (C) electrode implantation. (D) Post-operative MRI confirmed AAV injection success. Gd-contrast enhanced MRI intensity at the injection site, facilitating visualization. R: right hemisphere; L: left hemisphere. (E) Measurements of distances and volumes of Gd-enhanced regions. Viral suspension (20 μl) was injected either once (20) or three times (20×3). For the three-times injection, the viral suspension was delivered along the vertical trajectory: at the bottom of the target site, 3mm above, and 6mm above the initial injection point.

In the initial surgery, anatomical MRI identified the HPC as the target region (Figure 2A). Using trajectory planning based on the MPRAGE images (Figure 2B), we injected an AAV vector suspension mixed with Gd directly into the HPC. Post-operative MRI was performed to confirm the injection site and estimate the diffusion volume of the AAV suspension (Figure 2D). Following surgery, the pig underwent a recovery period of 8–12 weeks to ensure adequate opsin expression.

The second, non-survival surgery, focused on assessing the functionality of eOPN3-mediated optogenetic modulation (Figure 1). Anatomical MRI was again employed to localize the ANT and HPC, the key regions of interest in the Papez circuit (Figure 2A). After determining the target coordinates, fiber photometry was conducted to confirm eOPN3 expression within the HPC. An optic fiber was inserted along the planned trajectory, beginning approximately 30 mm from the target site. Fluorescence intensity increased progressively as the fiber approached the injection site, with a marked slope change indicating the optimal depth for implantation (Figure 1, bottom center).

Once photometry was complete, the optic fiber was replaced with a combined DBS electrode and optical fiber (Supplementary Figure 1B), which were bilaterally implanted into the ANT and HPC using Kopf stereotactic arms. In addition, another stereotactic arm equipped with an injection pump was positioned to allow pharmacological induction of epileptic activity.

To characterize the effects of optogenetic modulation, we recorded both electrical stimulation-evoked responses and LFPs simultaneously from the bilateral ANT and HPC under conditions with and without optical illumination. After completing the electrophysiological experiments, the pig was euthanized, and the brain was harvested for histological analysis to confirm the anatomical accuracy of the eOPN3 expression and to assess any associated neuropathological changes.

### Estimation of AAV Diffusion Using Gd-Contrast MRI

Determining the optimal volume of viral suspension required to achieve adequate coverage in the brain is a significant challenge. The diffusion and transduction of viral vectors are complex and influenced by multiple factors, including the specific brain subregion and its structural components, such as white matter fiber tracts and gray matter architecture, as well as properties of the viral injection, including volume, viral serotype, viscosity, and titer. To gain preliminary insights, our initial focus was on the volumetric measurement.

To facilitate this analysis, Gd was added to the viral suspension to enable visualization using MRI. Two different injection volumes were tested: 20 µl and 60 µl. For the 20 µl condition, a single injection was performed in the ANT. For the 60 µl condition, three sequential 20 µl injections were delivered along the longitudinal axis of the HPC. To control for tissue variability, in one pig (Pig 3), the left HPC received a single 20 µl injection, while the right HPC received three 20 µl injections at staggered depths (bottom of the target site, 3 mm above, and 6 mm above the initial injection point) (Figure 2B). This approach allowed for a direct comparison of diffusion under consistent tissue conditions.

Immediately following the AAV injection surgeries, post-operative MRI was performed using the same imaging sequences as the pre-operative scans. The inclusion of Gd in the viral suspension enhanced the MRI signal intensity, allowing for clear delineation of the diffusion area (Figure 2D). Using the MRI images, diffusion distances were measured along three axes: anterior-posterior (A-P), medial-lateral (M-L), and dorsal-ventral (D-V) (Figure 2E).

For statistical analysis, data from single 20 µl injections in the ANT (n=3) and HPC (n=1) were combined. There was no significant difference in diffusion distances between the 20 µl and 60 µl conditions along the A-P (5.6 ± 1.0 mm for 60 µl vs. 8.0 ± 0.4 mm for 20 µl, p=0.07) or M-L axes (5.2 ± 0.7 mm for 60 µl vs. 5.8 ± 0.4 mm for 20 µl, p=51). However, as expected, diffusion distances along the D-V axis were significantly greater for the 60 µl condition (13.2 ± 1.0 mm for 20 µl x3) compared to the 20 µl condition (9.3 ± 0.9 mm) (p=0.02; non-paired t-test with Welch’s correction). All values are reported as mean ± SEM.

### Optogenetic Modulation of Stimulation-Evoked Response Potentials (SERPs)

To investigate the effects of eOPN3-mediated optogenetic neuromodulation in the Papez circuit, LFPs and single-pulse SERPs were analyzed.

In the SERP experiment, two electrode contacts were used for stimulation, while the remaining contacts recorded the responses. The initial selection of stimulation contacts was guided by MRI-confirmed electrode placement, ensuring precise targeting. The final selection was refined based on the amplitude and waveform of recorded SERPs, optimizing signal fidelity. Single-pulse stimulation (6 mA, 250 μs pulse width, 0.2 Hz frequency) elicited SERPs, and stable baselines were established prior to the application of green light illumination (Figure 3). Green light was delivered continuously for 4–5 minutes at a specified power level (Figure 3, middle panel). To minimize potential artifacts and ensure accurate measurements, a 10-minute recovery period followed each illumination, allowing the baseline to be restored. Differential potentials between adjacent electrodes were analyzed, with the amplitude of the first post-stimulation peak quantified to assess eOPN3 modulation.

**Figure 3.**
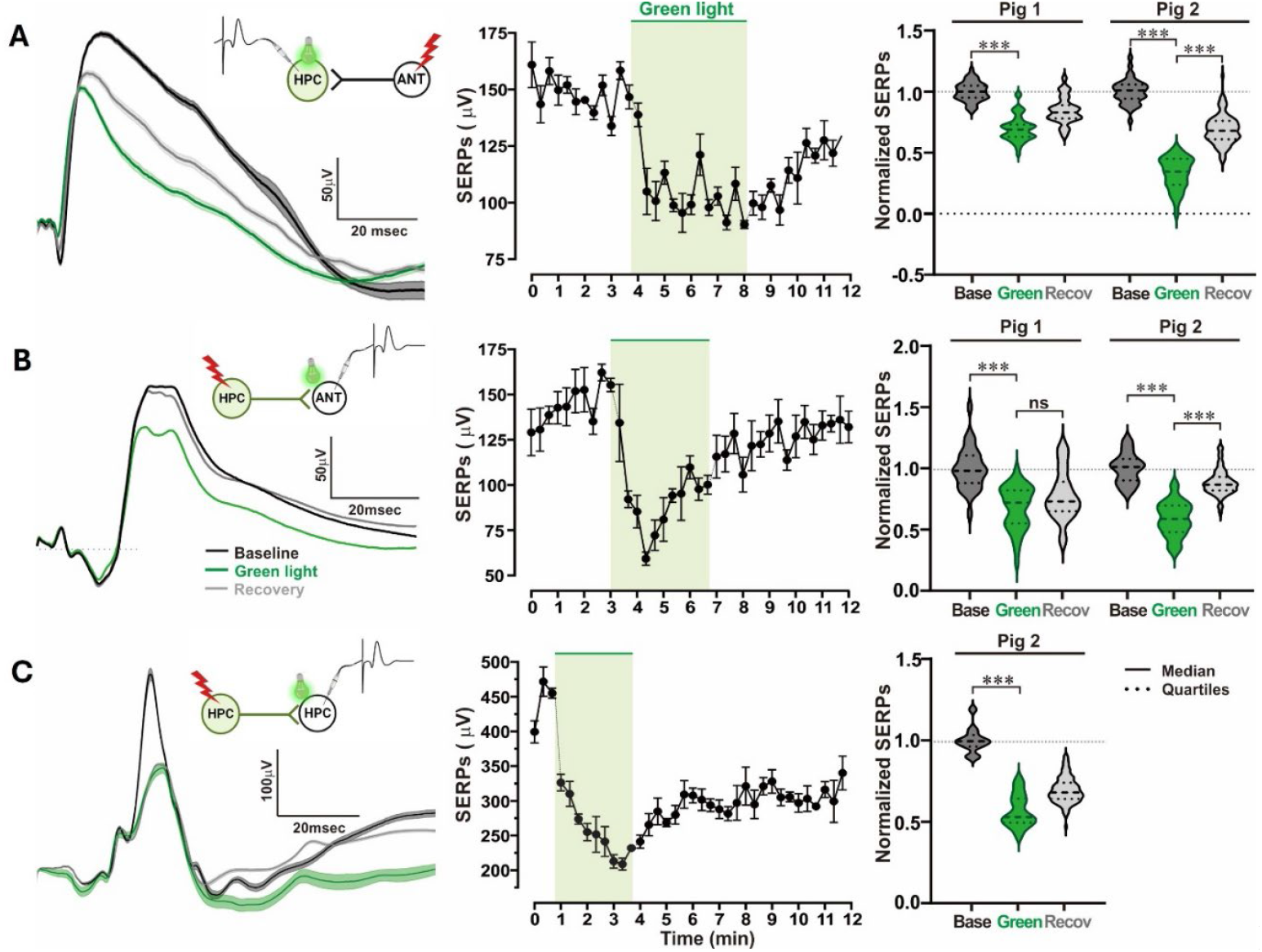
Optogenetic Modulation of Stimulation-Evoked Response Potentials (SERPs). The neuromodulatory effects of eOPN3 were characterized in the Papez circuit by stimulating and recording either the anterior nucleus of the thalamus (ANT) or hippocampus (HPC). (A) Soma Inhibition. AAV-CaMKII-eOPN3 was injected into the right HPC. Electrical stimulation of the ipsilateral ANT evoked robust synaptic responses in the HPC (black trace). Green light illumination significantly reduced SERPs (green trace), with recovery (gray trace) observed after light termination. The middle graph illustrates the temporal dynamics of light-induced SERP reduction, while box plots demonstrate subject-specific effects, showing differences between 8-week (Pig 1) and 12-week (Pig 2) expression durations. (B) Synaptic Inhibition: Scenario 1. Electrical stimulation was applied to the eOPN3-expressing HPC, with light illumination targeting the ipsilateral ANT to evaluate eOPN3 effects on synaptic release. Light illumination significantly reduced SERPs in the ANT. Temporal trajectories and subject-specific effects are also shown. (C) Synaptic Inhibition: Scenario 2. Illumination and recording were conducted in the contralateral HPC to assess eOPN3-mediated optogenetic effects on synaptic release. In the left panel, the averaged trace (solid line) from all raw SERP traces within each condition (before, during, and after illumination) and the standard error of the means (SEM, shadow) are presented. In the middle panel, to evaluate temporal patterns of SERPs responding to the illumination, SERPs’ amplitudes were averaged across four consecutive trials, with each data point representing this average. In the right panel, the amplitudes of SERPs were normalized by dividing the mean amplitude values of the baseline SERPs (Welch’s *t*-test, ***; p< 0.001).

Given the unilateral expression of eOPN3 in the HPC, three distinct modulatory paradigms were evaluated to elucidate its functional effects: somatic inhibition and two synaptic inhibition scenarios. First, somatic inhibition was characterized by illuminating the HPC while stimulating the ANT. Since single-pulse stimulation of the ANT primarily activates glutamatergic efferent fibers projecting to the HPC, we hypothesized that eOPN3-mediated potassium channel activation ^29^ would induce hyperpolarization, thereby reducing synaptic responses. As expected, ANT stimulation evoked robust HPC SERPs (black trace, Figure 3), which were significantly reduced during green light illumination (green trace). Notably, SERPs gradually recovered following light termination (gray trace), consistent with previous characterization of off-kinetics of eOPN3 ^29^ (Figure 3A). We performed Welch’s t-test to statistically compare the group means, accounting for unequal variances and sample sizes (***; *p* < 0.001). We then used a violin plot to visualize the groups’ distributions, which indicated potential differences in means and variances (right panels in Figure 3).

To further investigate eOPN3’s impact on synaptic release in HPC projections, single-pulse stimulation was applied to the eOPN3-expressing HPC, with SERPs recorded in the ipsilateral ANT or contralateral HPC. Importantly, illumination of the recording site significantly reduced SERPs in both the ipsilateral ANT (Figure 3B) and contralateral HPC (Figure 3C), indicating that eOPN3 modulation extended to synaptic terminals and highlighting its functional relevance across interconnected regions.

Finally, to examine the influence of viral expression duration on functional efficacy, two pigs received AAV-eOPN3 injections, with expression tests conducted at distinct time points: 8 weeks post-injection in Pig 1 and 12 weeks post-injection in Pig 2. Interestingly, in both subjects, SERPs were consistently reduced during light illumination (Figure 3A-C, right panels). Moreover, the magnitude of the light-induced effects was comparable across the two-time points, suggesting stable and effective opsin expression over time.

### Optogenetic Neuromodulation Effects on KA-Induced Seizures in the Hippocampus

Next, we evaluated the inhibitory effects of eOPN3 on KA-induced seizure-like activity. KA (1 µg/µl) was infused into the HPC using a needle attached to a Hamilton syringe. Neural activity was continuously recorded throughout the KA infusion and subsequent light illumination phases (Figure 4A). KA was delivered at a controlled rate of 0.2–1.0 µl/min using a micro pump, with a total of 1-2 µl infused per session. Following each infusion, a 10-minute observation period allowed seizure activity to develop, during which LFP recordings were performed via bilaterally implanted DBS electrodes in the ANT and HPC.

**Figure 4.**
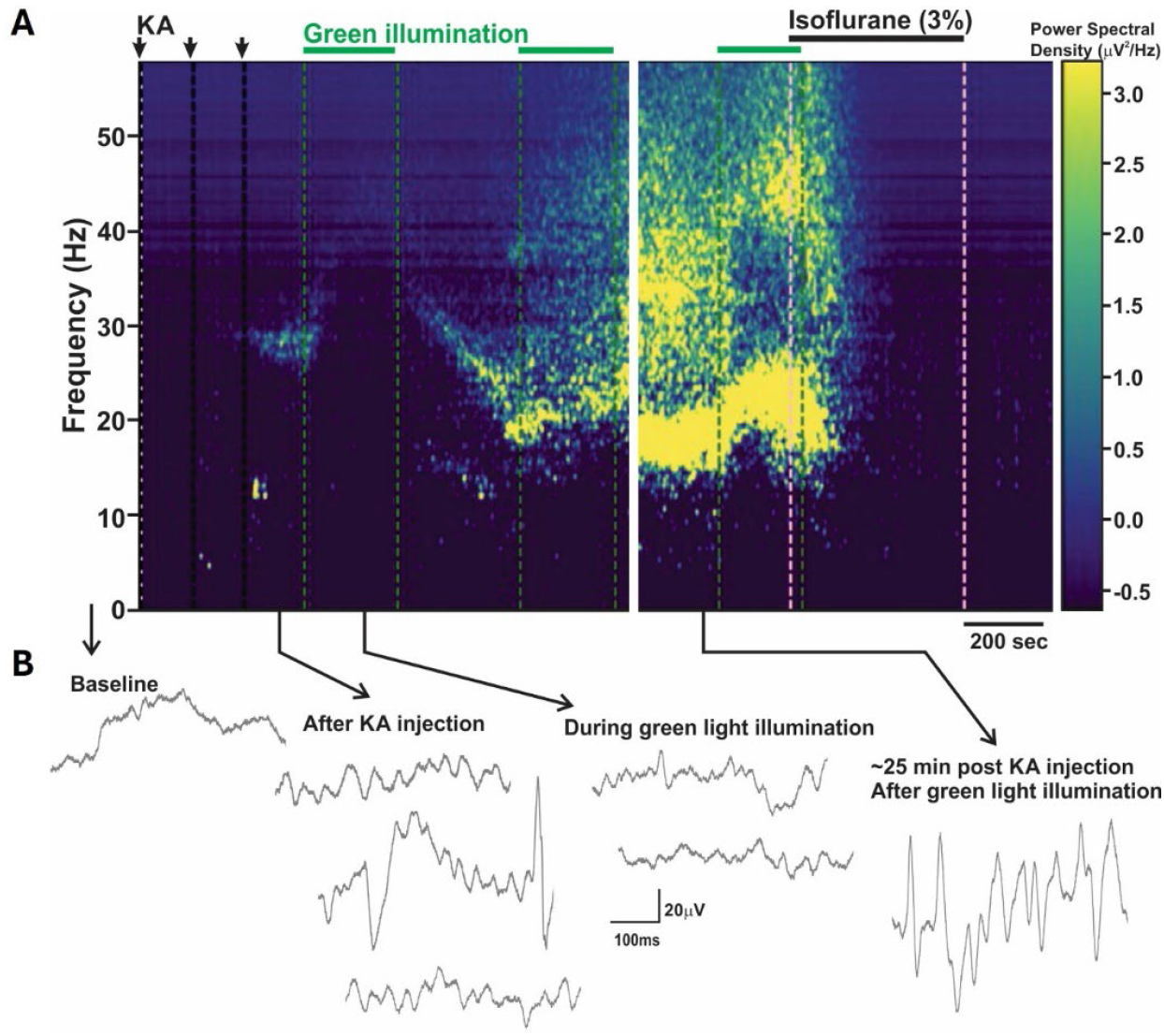
Optogenetic Effects on KA-Induced Seizure-Like Activity. (A) Spectrogram. Kainic acid (KA) injection into the HPC (arrows, 1 μl per injection) induced seizure-like activity characterized by 20-30 Hz oscillations. Early-stage KA-induced oscillations (∼30 Hz) were effectively suppressed by ANT-HPC illumination (green bars). (B) Representative hippocampal LFP traces from the KA-injection side, illustrating the modulation of seizure-like activity by optogenetic intervention.

Once seizure activity was visually confirmed, green light illumination was applied through all implanted optic fibers targeting the bilateral ANTs and HPCs (Figure 4A). The green light significantly attenuated KA-induced beta-band oscillations (∼30 Hz) in the intrahippocampal region. However, once KA-induced generalized oscillatory activity had propagated to the contralateral HPC and the ANT, the optogenetic modulation was unable to suppress the ongoing seizure-like activity. Representative LFP traces illustrating these findings are shown in Figure 4B.

### Expression of eOPN3 in the Hippocampus

After the electrophysiology recording completion, the brain was extracted and later processed for histological assessment. Representative images were obtained from Pig 3, which received 20 µl and 60 µl viral suspension injections in the left and right HPCs, respectively. Brightfield images (Figure 5A) were annotated (1 to 5) to compare HPC structures and positions along the A-P axis. These images were sequentially arranged from the anterior (A) to posterior (P) regions, followed by corresponding fluorescent images (Figure 5B and C) for detailed analysis.At anterior levels (Levels 1–3), eOPN3 expression was detected in both the left and right HPCs (Figure 5B). However, at more posterior levels (Levels 4–5), eOPN3 expression was predominantly localized in the right HPC, which corresponded to the higher injection volume (Figure 5B, 4R and 5R). Cellular level eOPN3 expression was clearly visualized in Figure 5C.

**Figure 5.**
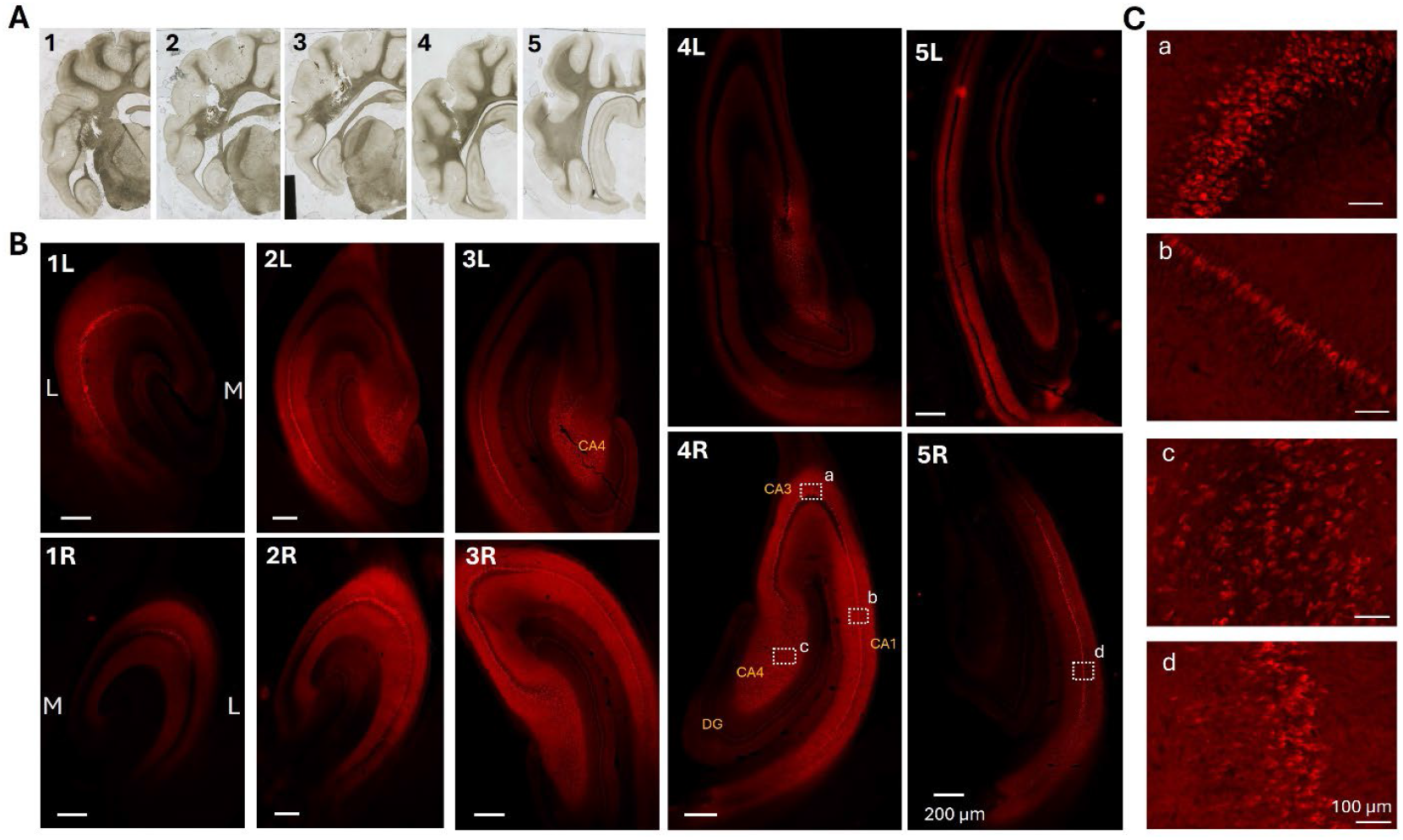
Histological Confirmation of eOPN3 Expression. (A) Bright-field images of pig brain slices. Numerical labels (1 through 5) indicate anterior-to-posterior levels in both (A) and (B). (B) Fluorescent images showing eOPN3 expression corresponding to the levels indicated in (A). The letter ‘L’ in the top-left corner of fluorescent images refers to the left hemisphere, and ‘R’ refers to the right hemisphere. One 20 μl AAV injection was made into the left HPC, while three injections (20 μl each) were performed in the right HPC. M and L in the middle of the images denote medial and lateral directions, respectively. (C) Cellular-level expression of eOPN3. The regions (a)-(d) are identified in slices 4R and 5R in panel (B).

These findings suggest that larger injection volumes result in more extensive eOPN3 expression, both in the number of expressing cells and the spatial distribution of expression. Notably, some slices exhibited strong eOPN3 expression in regions without apparent cell body expression, which likely reflects eOPN3 expression in axonal and other cellular compartments and suggests that functional effects of eOPN3 extend beyond the immediate injection sites. This widespread expression pattern highlights the potential of eOPN3 to modulate neuronal activity across broader regions of the HPCus, consistent with its design for enhanced membrane targeting and function.

## Discussion

Optogenetics represents a potentially groundbreaking approach in the treatment of epilepsy, leveraging cell-type-specific control of neuronal activity via light-sensitive opsins ^32^. This technology enables the precise modulation of excitatory or inhibitory neuronal populations, offering the potential to modulated brain excitability and suppress seizures without impairing normal brain function. As the field of neuromodulation evolves, the role of optogenetics is poised to expand significantly. In this study, we establish the feasibility of a porcine seizure model as a robust translational platform for optogenetic neuromodulation in TLE. Using eOPN3, a highly sensitive and bistable inhibitory opsin, we demonstrated that targeted modulation of HPC and ANT activity has a strong potential to effectively suppress Papez circuit synaptic activity and pathological seizure-like neural activity, underscoring the therapeutic promise of optogenetic interventions for neurological diseases.

eOPN3 addresses many limitations of earlier opsins, offering distinct advantages for clinical translation. Its high sensitivity to low-intensity red light (∼5–20 µW/mm^2^) minimizes tissue heating, facilitating safer applications in larger brains. Testing in the porcine brain confirmed that the inhibitory effects of red illumination were comparable to those achieved with green light (data not shown). However, the deeper penetration and slower kinetics of red light limited the extent of our testing within the constraints of experimental timelines. Nonetheless, the red-shifted action spectrum enhances tissue penetration and reduces light scattering, further establishing its suitability for clinical applications. eOPN3 also leverages the G_i/o_ signaling pathway, which suppresses calcium influx and inhibit the presynaptic SNARE complex to achieve long-lasting inhibition of synaptic transmission with a single light pulse ^33,34^. Its bistable property eliminates the need for continuous illumination, reducing the risk of tissue damage and enhancing the efficiency of neuromodulation. These attributes make eOPN3 a precise, long-lasting, and minimally invasive tool for controlling epileptic networks.

In our study, the CaMKII promoter enabled cell-type-specific expression of eOPN3, predominantly in pyramidal neurons within the CA1, CA3, and CA4 regions (Figure 5). Notably, eOPN3 expression in the granule cell layer of the DG was comparatively low (Figure 5). While granule cells do express CaMKII, their expression levels could be lower than those in pyramidal neurons ^35^. Since KA strongly activates granule cells in the DG due to the high prevalence of KA receptors ^36,37^, the relatively low eOPN3 expression in this region may diminish its inhibitory efficacy on seizure activity. This finding underscores the importance of considering seizure type, receptor distribution, and promoter selection to optimize optogenetic seizure control. Future studies should explore these factors in scalable *in vivo* systems to further refine therapeutic outcomes.

Several key translational challenges remain to be addressed, including optimizing vector injection volume, injection site distribution, opsin longevity and safety, and the relationship between injection parameters and viral expression. Understanding the underlying network mechanisms and identifying precise therapeutic targets will require advanced in vivo studies in large animal models. Additionally, refining implantable hardware and delivery techniques will be critical to advancing optogenetic therapies toward clinical use. The porcine brain provides an ideal experimental platform for these investigations due to its larger size and cortical complexity, which better emulate human neuroanatomy.

This study provides a foundation for translating optogenetic neuromodulation into clinical practice, highlighting the potential of eOPN3 as a novel treatment for drug-resistant epilepsy and providing a large animal *in vivo* model. By demonstrating the efficacy of eOPN3 in modulating seizure activity within a large animal model, we bridge a critical gap between rodent studies and human clinical applications. Future investigations should focus on optimizing this platform for epilepsy and extending its applicability to other neurological disorders characterized by network hyperexcitability, paving the way for broader clinical adoption of optogenetic therapies.

## Acknowledgment

This research was supported by NIH (R01NS092882), Birdel Fund Mayo Clinic Philanthropy and Cadence Neuroscience who provided the electrophysiology neural workstation and the INSR devices. We thank the XRI Core and Department of Comparative Medicine within Mayo Clinic and Dr. Jean-Charles Paterna, the head of the UZH viral vector core, for supporting our experiments.

Acknowledgment

This research was supported by Birdel Fund Mayo Clinic Philanthropy and Cadence Neuroscience who provided the electrophysiology neural workstation and the INSR devices. We thank the XRI Core and Department of Comparative Medicine within Mayo Clinic and Dr. Jean-Charles Paterna, the head of the UZH viral vector core, for supporting our experiments.

## Figures

**Supplementary Figure 1.**
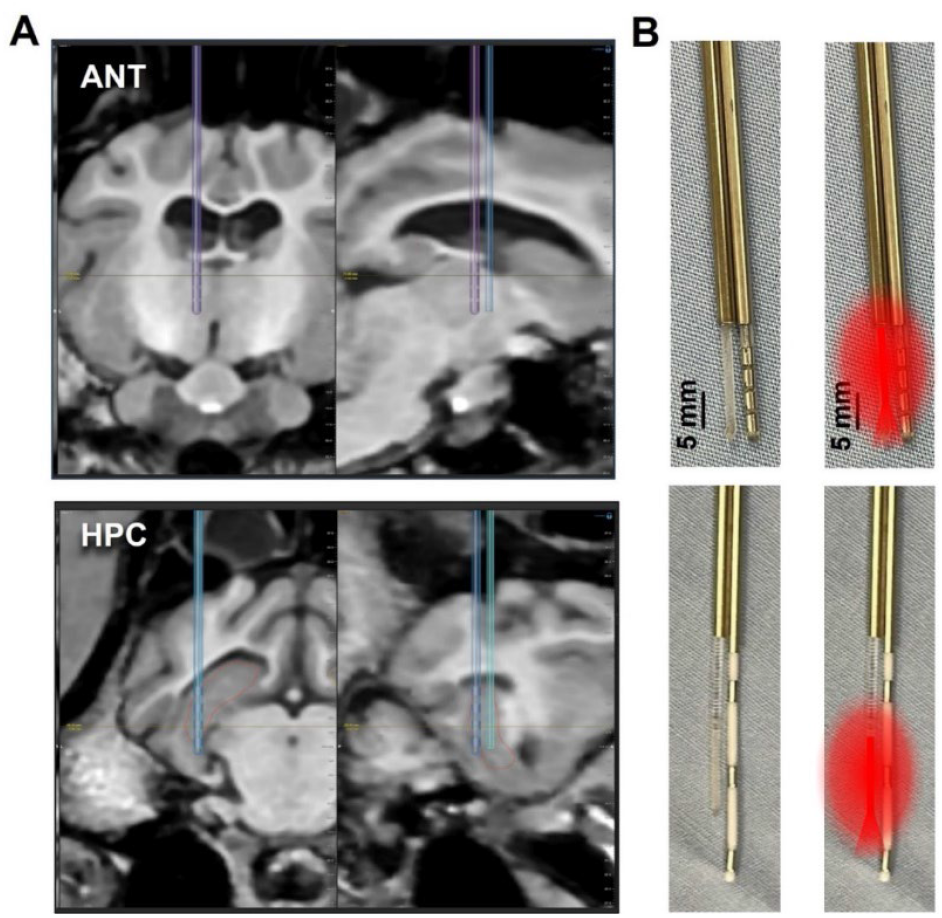
MRI Planning for DBS Electrode Implantation and Design of Implantable Electrodes. (A) Coronal and sagittal MRI images targeting the ANT (top) and HPC (bottom). These images were used to plan precise electrode trajectories. (B) The design and placement of DBS electrodes combined with an optic fiber are shown. Top, in the ANT and HPC, and bottom, in the HPC. Right panels show an illustration of light illumination profile around the filled 10mm at the tip of the fiber.

**Supplementary Figure 2.**
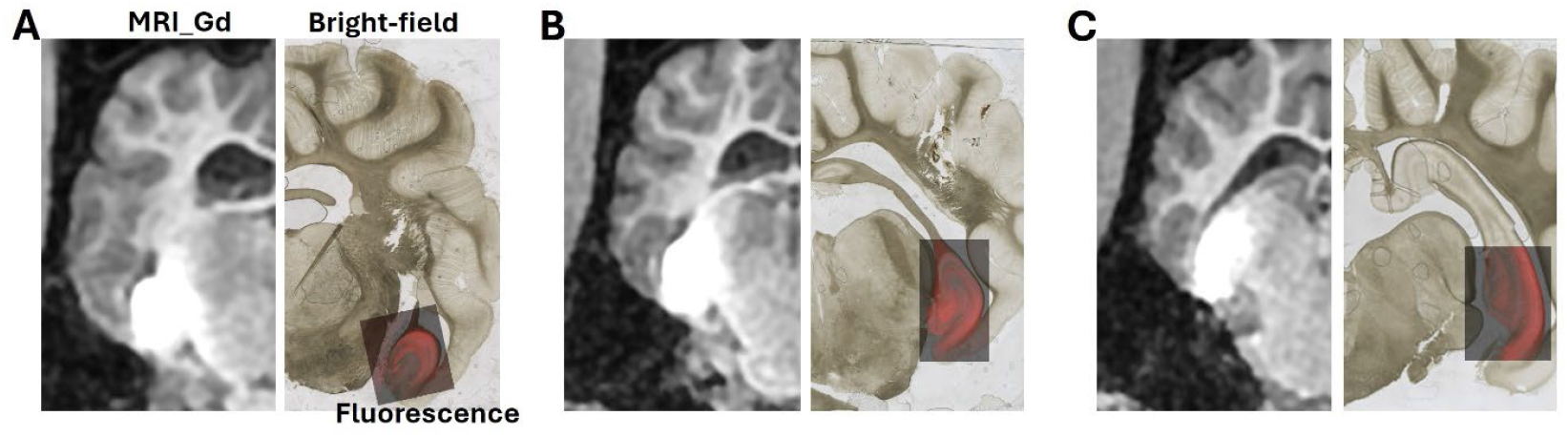
Combined Gd-Enhanced MRI, Bright-Field, and Fluorescent Images of the Hippocampus. Three levels (A, B, and C) were selected from the anterior-to-posterior series (1, 3, and 5, respectively) in Figure 5 to correlate Gd-enhanced MRI with histological data. For each selected level, the figure displays (1) Gd-enhanced MRI to identify the injection site and diffusion pattern, (2) bright-field images for anatomical landmark identification, and (3) overlayed fluorescent images to highlight eOPN3 expression in the HPC. These combined images provide spatial context for the HPC, Gd distribution, and eOPN3 expression, facilitating anatomical and functional correlation.

